# WDR2 regulates the orphan kinesin KIN-G to promote hook complex and Golgi biogenesis in *Trypanosoma brucei*

**DOI:** 10.1101/2025.01.29.635568

**Authors:** Qing Zhou, Huiqing Hu, Ziyin Li

## Abstract

The flagellum in *Trypanosoma brucei* plays crucial roles in cell locomotion, cell morphogenesis, and cell division, and its inheritance depends on the faithful duplication of multiple flagellum-associated structures. One of such cytoskeletal structures is a hairpin-like structure termed the hook complex composed of a fishhook-like structure and a centrin arm structure, whose cellular functions remain poorly understood. We recently identified KIN-G, an orphan kinesin that promotes hook complex and Golgi biogenesis. Here we report a WD40 repeats-containing protein named WDR2, which interacts with and regulates KIN-G. WDR2 co-localizes with KIN-G at the centrin arm, and knockdown of WDR2 disrupts hook complex integrity and morphology, inhibits flagellum attachment zone elongation and flagellum positioning, and eventually arrests cytokinesis. Knockdown of WDR2 also disrupts the maturation of the centrin arm-associated Golgi, thereby impairing Golgi biogenesis. WDR2 interacts with KIN-G via its N-terminal unknown motifs, the middle domain containing a coiled coil and a PB1 motif, and the C-terminal WD40 domain, and targets KIN-G to its subcellular location. These results uncover a regulatory role for WDR2 in recruiting KIN-G to regulate hook complex and Golgi biogenesis, thereby impacting flagellum inheritance and cell division plane positioning.

## Introduction

The flagellum in the parasitic protozoan *Trypanosoma brucei*, the causative agent of sleeping sickness in humans and nagana in livestock in sub-Saharan Africa, plays essential roles in cell motility, cell division, and cell-cell communication. It is nucleated from a centriole-like structure termed the basal body, and it exits the cell through the flagellar pocket and attaches to the cell membrane for most of its length via a specialized cytoskeletal structure termed the flagellum attachment zone (FAZ) (Sunter and Gull, 2016), which maintains flagellum attachment and defines the cell division plane (Vaughan et al., 2008; Zhou et al., 2011). In the flagellar pocket region, several specialized cytoskeletal structures, including the flagellar pocket collar (FPC) (Lacomble et al., 2009) and the hook complex (Esson et al., 2012), associate with the flagellum (Fig. 1A). The FPC is a horseshoe-shaped structure that wraps around the flagellum and likely plays roles in restricting protein transport in and out of the cell through the flagellar pocket (Lacomble et al., 2009). Sitting at the top of the FPC (Esson et al., 2012), the hook complex is a hairpin-like structure composed of a fishhook-like structure marked by TbMORN1 and TbLRRP1 (Morriswood et al., 2009; Zhou et al., 2010) and a bar-shaped centrin arm structure marked by two centrin proteins TbCentrin2 and TbCentrin4 (He et al., 2005; Shi et al., 2008). The centrin arm sits alongside the shank part of the fishhook-like structure, with the centrin arm and the shank embedding the proximal end of the intracellular FAZ filament (Esson et al., 2012; Morriswood, 2015) (Fig. 1A). The structural integrity of the hook complex is critical for promoting FAZ elongation and flagellum positioning, thereby impacting the placement of the cell division plane and cytokinesis (Pham et al., 2020; Zhou et al., 2010).

**Figure 1.**
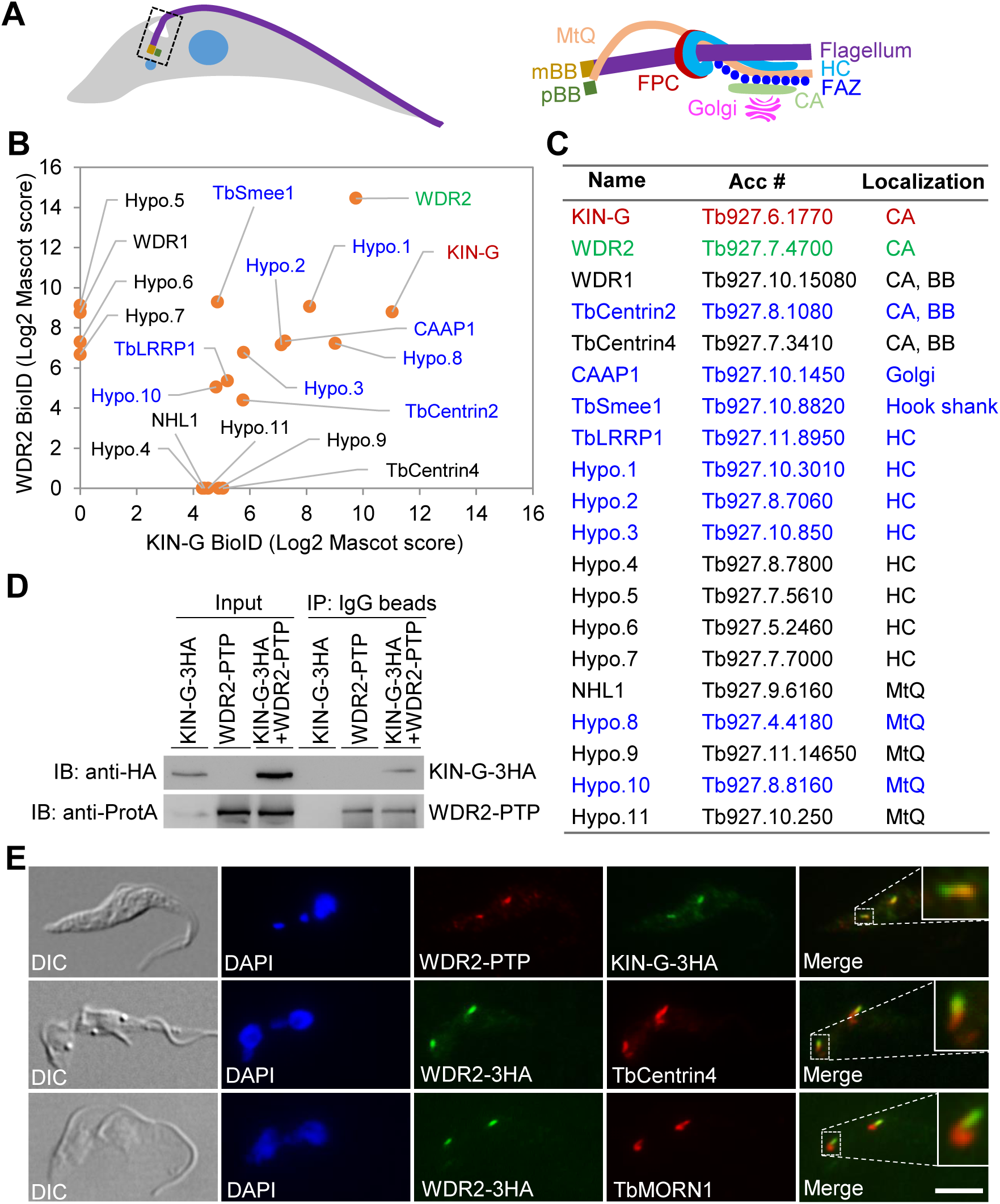
Identification of WDR2 as an interacting protein of KIN-G. (**A**). Schematic drawing of a trypanosome cell showing the flagellum and its associated cytoskeletal structures. mBB: mature basal body; pBB: pro-basal body; FPC: flagellar pocket collar; MtQ: microtubule quartet; FAZ: flagellum attachment zone; HC: hook complex; CA: centrin arm. (**B**). KIN-G- and WDR2-proximal proteins identified by BioID. Bait proteins are highlighted in red or green, and common proximal proteins are highlighted in blue. (**C**). List of KIN-G- and WDR2-proximal proteins and their subcellular localizations. Bait proteins are in red or green, and KIN-G- and WDR2-shared proximal proteins are in blue. CA: centrin arm; BB: basal body; HC: hook complex; MtQ: microtubule quartet. (**D**). Co-immunoprecipitation to test the *in vivo* interaction between KIN-G and WDR2. Endogenous KIN-G-3HA and WDR2-PTP were detected by anti-HA and anti-Protein A antibodies, respectively. (**E**). Subcellular localization of WDR2, endogenously tagged with PTP or 3HA, and its co-localization with endogenously 3HA-tagged KIN-G, TbCentrin4, and TbMORN1. Insets show the zoom-in view of the selected region. Scale bar: 5 μm.

Alongside the intracellular FAZ filament runs a specialized set of four microtubules termed the microtubule quartet (MtQ) (Fig. 1A), which originates between the mature and the pro-basal bodies, wraps around the flagellar pocket, and then pass through the FPC and the hook complex to extend to the distal cell tip (Esson et al., 2012; Vaughan and Gull, 2016). The precise molecular function of the MtQ remains elusive, but three MtQ proximal end-localized proteins, TbSpef1, NHL1, and SNAP1, appear to promote basal body rotation and segregation, thereby facilitating flagellum positioning (Gheiratmand et al., 2013; Pham et al., 2022; Souza Onofre et al., 2023). In close proximity to the centrin arm, there sits the Golgi apparatus (Fig. 1A), which is duplicated through a *de novo* pathway, by which the new Golgi is assembled next to the old Golgi and the ER exit site (He et al., 2004). Duplication of the single Golgi apparatus in the procyclic (insect) form of *T. brucei* appears to depend on the assembly of an intact centrin arm, as knockdown of TbCentrin2 (He et al., 2005) and the centrin arm-localized TbPLK (*T. brucei* Polo-like kinase) (de Graffenried et al., 2008) disrupted Golgi biogenesis. Although knockdown of TbCentrin2 and TbPLK exerts distinct effects on Golgi biogenesis, with TbCentrin2 depletion completely inhibiting the formation of new Golgi (He et al., 2005) and TbPLK depletion producing numerous small Golgi (de Graffenried et al., 2008), the results suggest that the centrin arm determines the Golgi assembly site. We recently identified and characterized an orphan kinesin named KIN-G, which localizes to the centrin arm and regulates hook complex and Golgi biogenesis in procyclic trypanosomes (Zhou et al., 2024). We also identified CAAP1 as a Golgi peripheral protein, which promotes the association between Golgi and the centrin arm to facilitate Golgi biogenesis (Zhou et al., 2024). KIN-G is a microtubule plus end-directed motor protein; hence, it might transport certain protein and/or non-protein cargos for hook complex and Golgi biogenesis. The cargos that KIN-G transports to the centrin arm and/or the Golgi remain unidentified.

In the current work, we attempted to identify the interacting proteins and the potential cargo proteins of KIN-G by proximity-dependent biotin identification (BioID). We identified a KIN-G-interacting protein named WDR2, a kinetoplastid-specific protein containing a WD40-repeats domain at the C-terminus, two domains of unknown function (DUF) at the N-terminus, and a small coiled-coil motif and a PB1 (Phox and Bem1) motif in the middle region, and showed that WDR2 plays an essential role in promoting hook complex and Golgi biogenesis in the procyclic form of *T. brucei*. We further demonstrated that WDR2 executes its function by recruiting and maintaining KIN-G at the centrin arm. These results discovered a new protein implicated in hook complex and Golgi biogenesis in *T. brucei* and uncovered its mechanistic role in the regulation of KIN-G localization and stability.

## Results

### Identification of WDR2 as a KIN-G-interacting protein localized at the centrin arm

We recently reported the essential role of the orphan kinesin KIN-G in regulating hook complex assembly and Golgi biogenesis in procyclic trypanosomes (Zhou et al., 2024). To identify the interacting partner(s) of KIN-G and the potential cargo proteins that KIN-G transports, we carried out BioID using KIN-G as bait. The KIN-G-proximal proteins thus identified were screened based on their subcellular locations that were determined previously by the TrypTag project (Billington et al., 2023). Since KIN-G localizes to the centrin arm (Zhou et al., 2024), only those proteins that localize to the close proximity of the centrin arm were considered as KIN-G-proximal proteins, which included the proteins localized to the centrin arm, the hook complex, and the MtQ (Fig. 1B). Among the KIN-G-proximal proteins, a WD40 repeats-containing protein (accession number: Tb927.7.4700) was the top hit and, thus, was chosen for further characterization. We named this protein WDR2 for <u>WD</u>40 <u>R</u>epeats-containing protein <u>2</u>, following the previously characterized WDR1 (Hu et al., 2017). Further, we performed BioID with WDR2 as bait, and the WDR2-proximal proteins were similarly screened based on their previously determined subcellular localizations, and were plotted, based on the mass spectrometry Mascot score, together with the KIN-G-proximal proteins to identify common proximal proteins (Fig. 1B). The list of proximal proteins of KIN-G and WDR2 included three centrin arm- and basal body-localized proteins, eight hook complex-localized proteins, five MtQ proximal end-localized proteins, one hook shank-localized protein (TbSmee1), and one Golgi-localized protein (CAAP1) (Fig. 1C). Other potential proximal proteins, including FAZ proteins, FPC proteins, and cytoskeletal proteins, were also identified (Supplemental Table S1). However, these proteins were not considered as potential interacting partners of KIN-G and/or WDR2 due to their farther distance to the centrin arm. Nine proximal proteins were identified by both KIN-G BioID and WDR2 BioID (Fig. 1B, C, protein names in blue), including TbCentrin2, CAAP1, TbSmee1, four hook complex-localized proteins, and two MtQ proximal end-localized proteins (Fig. 1B, C).

To test whether WDR2 interacts with KIN-G *in vivo* in trypanosomes, we epitope-tagged KIN-G and WDR2 at their respective endogenous locus in the same cell line and then carried out co-immunoprecipitation. Immunoprecipitation of WDR2-PTP was able to pull down KIN-G-3HA from the cell lysate (Fig. 1D), confirming that the two proteins form a complex in trypanosomes. Further, co-immunostaining showed that WDR2 and KIN-G co-localized, and co-immunostaining with the centrin arm marker TbCentrin4 and the hook complex marker TbMORN1 confirmed the localization of WDR2 to the centrin arm, alongside the shank part of the hook complex (Fig. 1E). Therefore, WDR2 and KIN-G form a complex and may function together at the centrin arm.

### Knockdown of WDR2 causes defective cytokinesis and FAZ elongation

To investigate the function of WDR2 in *T. brucei*, we performed RNAi in the procyclic form. Western blotting was first carried out to monitor the protein level of WDR2, which was endogenously tagged with a triple HA epitope, before and after WDR2 RNAi induction. WDR2 protein level was significantly reduced after one day and was undetectable at day 3 of RNAi induction (Fig. 2A). The depletion of WDR2 by RNAi caused severe growth defects and eventual cell death (Fig. 2B), suggesting the essentiality of WDR2 for cell viability in procyclic trypanosomes. Knockdown of WDR2 caused a substantial increase of xNyK (x>2, y≥1) cells and the emergence of abnormal cell types, such as 2N1K cells and 0N1K cells (Fig. 2C), which could be derived from the aberrant division of 2N2K cells. These results demonstrated that WDR2 knockdown impaired cytokinesis, although this could be the secondary effects attributed to the defects in FAZ elongation and flagellum positioning (see Fig. 2G-I below). Since the earliest time point for WDR2 RNAi to generate growth defects and cytokinesis defects was 48 h, all the subsequent phenotypic analyses were carried out with cells induced for WDR2 RNAi for 48 h.

**Figure 2.**
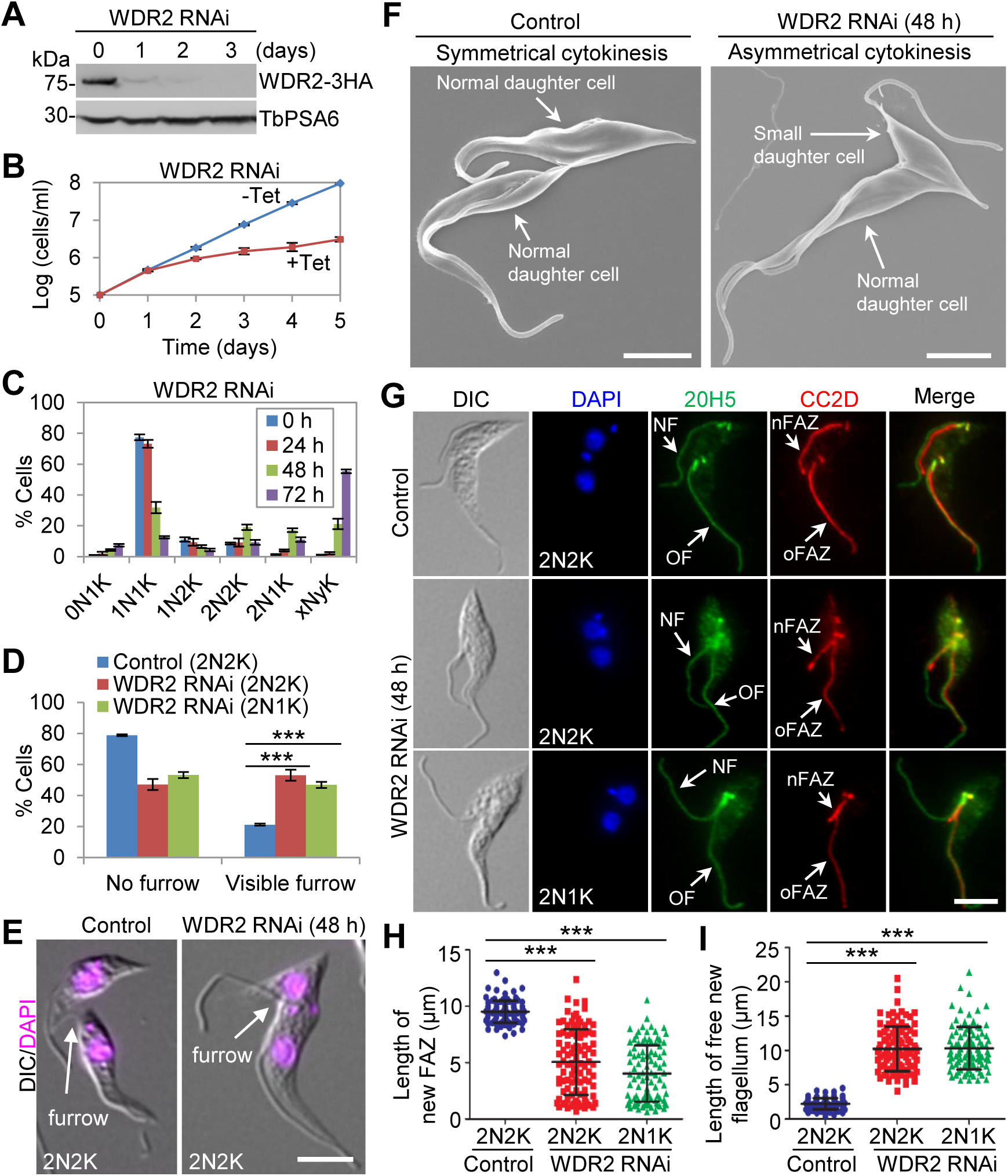
Knockdown of WDR2 causes asymmetrical cytokinesis and inhibits FAZ elongation. (**A**). Knockdown of WDR2 by RNAi. Endogenous WDR2-3HA was detected by the anti-HA antibody. TbPSA6 served as a loading control. (**B**). WDR2 knockdown caused growth defects in procyclic trypanosomes. (**C**). Effect of WDR2 knockdown on cell cycle progression. Shown is the quantitation of cells with different numbers of nuclei (N) and kinetoplasts (K) before and after tetracycline (Tet)-induced WDR2 RNAi induction. Error bars indicate S.D. from three independent experiments. (**D**). Effect of WDR2 knockdown on cleavage furrow ingression. Shown is the percentage of bi-nucleated cells with or without a visible cleavage furrow. Error bars indicate S.D. from three independent experiments. ***, *p*<0.001. (**E**). Control and WDR2 knockdown cells undergoing cytokinesis. Scale bar: 5 μm. (**F**). Scanning electron microscopic analysis of control and WDR2 RNAi cells undergoing cytokinesis. Scale bar: 5 μm. (**G**). WDR2 knockdown disrupted FAZ elongation and flagellum positioning. Cells were co-immunostained with 20H5, which labels the flagellum, and anti-CC2D antibody, which labels the FAZ filament. NF: new flagellum; OF: old flagellum; nFAZ: new FAZ; oFAZ: old FAZ. Scale bar: 5 μm. (**H**). Measurement of the length of the new FAZ in bi-nucleated cells before and after WDR2 RNAi. ***, *p*<0.001. (**I**). Measurement of the length of the free new flagellum in bi-nucleated cells before and after WDR2 RNAi. ***, *p*<0.001.

After WDR2 RNAi, the bi-nucleated (2N2K and 2N1K) cells with a visible cleavage furrow were significantly increased (Fig. 2D). Moreover, in the WDR2 RNAi-induced cell population, almost all of the dividing bi-nucleated cells appeared to produce a smaller-sized daughter cell containing a long unattached flagellum and a normal-sized daughter cell containing a normally attached flagellum, in striking contrast to that in control cells where the two daughter cells had normally attached flagella and similar cell size (Fig. 2E, F). In wild-type trypanosomes, the two daughters of a dividing cell are not identical in size (Abeywickrema et al., 2019), but the new-flagellum daughter cell of the dividing WDR2 RNAi cell was much smaller than that of the dividing control cell (Fig. 2E, F). Thus, knockdown of WDR2 caused asymmetrical cytokinesis, likely due to cell division plane misplacement.

The production of smaller-sized new flagellum daughter cells after WDR2 RNAi prompted us to examine the effect of WDR2 RNAi on FAZ assembly and flagellum attachment by immunofluorescence microscopy using the anti-CC2D antibody, which labels the intracellular FAZ filament (Zhou et al., 2011), and the pan-centrin antibody 20H5, which labels the flagellum in addition to the basal body and the centrin arm (He et al., 2005). The results showed that WDR2 knockdown produced bi-nucleated cells with a shorter new FAZ and a longer unattached new flagellum (Fig. 2G-I). Measurement of the length of the new FAZ and the length of the unattached (free) new flagellum showed that the average length of the new FAZ in 2N2K cells and 2N1K cells was significantly reduced after WDR2 RNAi induction (Fig. 2H). Consequently, the average length of the unattached new flagellum or free new flagellum in 2N2K cells and 2N1K cells was significantly increased after WDR2 RNAi (Fig. 2I). These results suggest that WDR2 knockdown disrupted the elongation of the new FAZ. Because the length of the new FAZ determines the position of the cell division plane in procyclic trypanosomes (Zhou et al., 2011) and WDR2 RNAi disrupted new FAZ elongation (Fig. 2G, H), the aberrant cytokinesis caused by WDR2 RNAi (Fig. 2E, F) was attributed to the disruption in FAZ elongation, which impaired the placement of the cell division plane.

### Knockdown of WDR2 disrupts flagellum positioning

Since the elongation of the new FAZ is coordinated with the migration of the new flagellum toward cell posterior (Sunter and Gull, 2016), the defective FAZ elongation caused by WDR2 RNAi may inhibit flagellum positioning. Indeed, immunostaining of WDR2 RNAi cells with the 20H5 antibody showed that the base of the new flagellum of the dividing bi-nucleated cells was positioned in close proximity to that of the old flagellum (Fig. 2F). This observation was confirmed by scanning electron microscopy, which showed that the new flagellum was positioned either closer to or in the proximity of the old flagellum in the dividing WDR2 RNAi cells (Fig. 3A). These results suggest that WDR2 depletion caused defective segregation and positioning of the new flagellum. Further, we investigated the effect of WDR2 RNAi on the segregation of the basal body, which nucleates the flagellum (Hu et al., 2015a), by immunofluorescence microscopy with the anti-TbSAS-6 antibody that detects the basal body cartwheel protein SAS-6 (Hu et al., 2015a). After WDR2 RNAi, the average inter-basal body distance in bi-nucleated cells was significantly reduced in 2N2K and 2N1K cells (Fig. 3B, C). Finally, we investigated the effect of WDR2 RNAi on the segregation of flagellum-associated structures, such as the centrin arm and the FPC by immunofluorescence microscopy with the 20H5 antibody and the anti-TbBILBO1 antibody, respectively (Fig. 3D-G). The results showed that the average inter-centrin arm distance and the average inter-FPC distance were both significantly reduced (Fig. 3D-G). These results provided further evidence that WDR2 depletion disrupted flagellum positioning.

**Figure 3.**
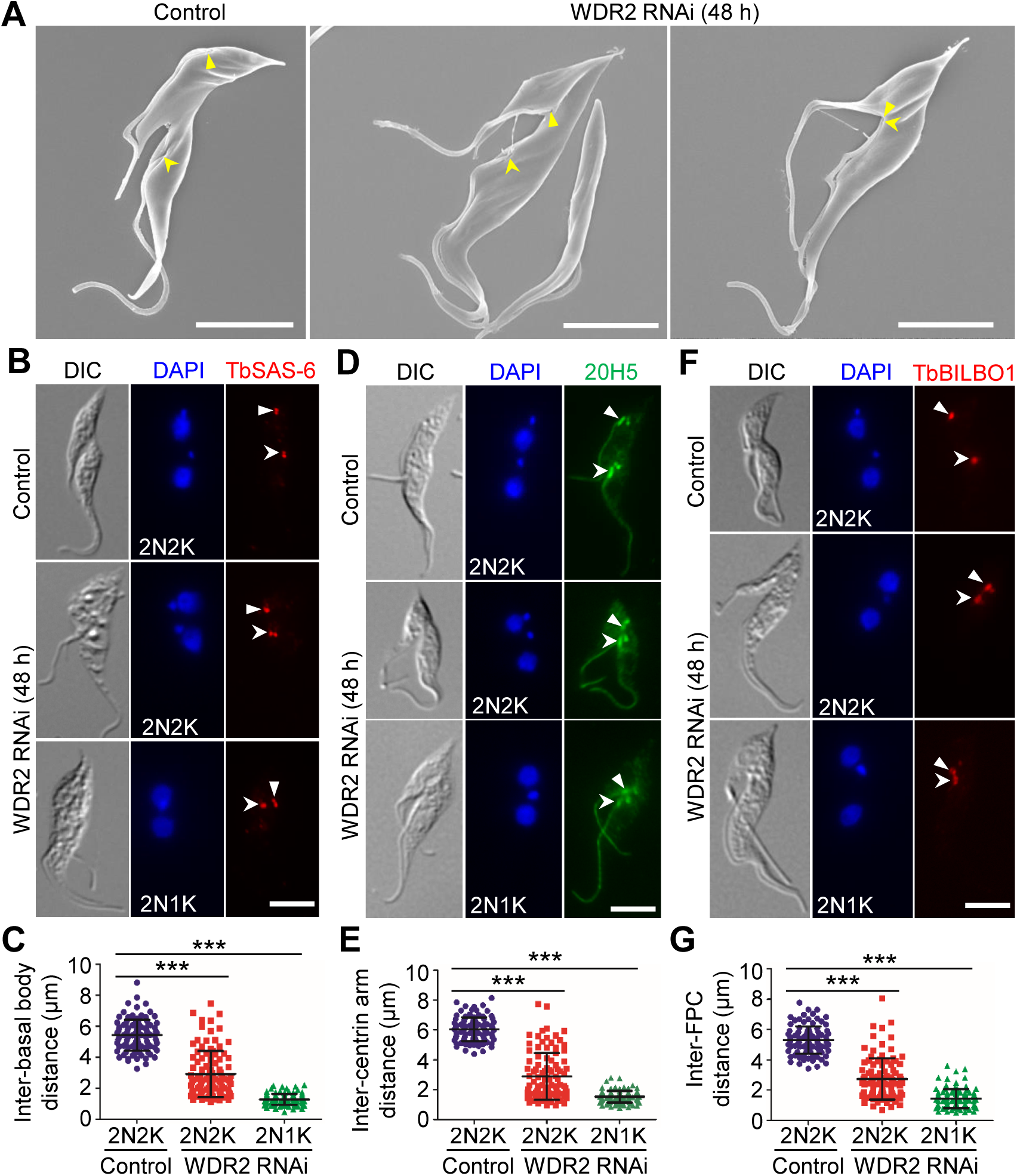
Knockdown of WDR2 disrupts flagellum positioning. (**A**). Scanning electron microscopic analysis of the dividing control and WDR2 RNAi cells, showing the effect of WDR2 knockdown on flagellum positioning. *Solid arrowheads* and *open arrowheads* indicate the proximal bases of the new and the old flagella, respectively. Scale bar: 5 μm. (**B**). Immunofluorescence microscopy to detect the basal body in control and WDR2 RNAi cells with the anti-TbSAS-6 antibody. *Solid arrowheads* and *open arrowheads* indicate the new and the old basal bodies, respectively. Scale bar: 5 μm. (**C**). Measurement of the inter-basal body distance in bi-nucleated cells before and after WDR2 RNAi. ***, *p*<0.001. (**D**). Immunofluorescence microscopy to detect the centrin arm in control and WDR2 RNAi cells with the pan-centrin antibody 20H5. *Solid arrowheads* and *open arrowheads* indicate the new and the old centrin arms, respectively. Scale bar: 5 μm. (**E**). Measurement of the inter-centrin arm distance in bi-nucleated cells before and after WDR2 RNAi. ***, *p*<0.001. (**F**). Immunofluorescence microscopy to detect the PFC in control and WDR2 RNAi cells with the anti-TbBILBO1 antibody. *Solid arrowheads* and *open arrowheads* indicate the new and the old FPCs, respectively. Scale bar: 5 μm. (**G**). Measurement of the inter-PFC distance in bi-nucleated cells before and after WDR2 RNAi (48 h). ***, *p*<0.001.

### Knockdown of WDR2 disrupts hook complex morphology and integrity

Since WDR2 localizes to the centrin arm and forms a complex with KIN-G, which is required for hook complex assembly (Zhou et al., 2024), we asked whether WDR2 plays similar roles. We first performed co-immunofluorescence microscopy using the pan-centrin antibody 20H5 and the anti-TbCentrin4 antibody and examined the integrity of the centrin arm in bi-nucleated cells before and after WDR2 RNAi induction. While the average length of the old centrin arm was not significantly affected by WDR2 RNAi, the average length of the new centrin arm was reduced by ∼43% from ∼1.3µm to ∼0.7µm after RNAi induction (Fig. 4A, B). Those bi-nucleated cells with the new centrin arm longer than 1.0µm were significantly reduced, whereas those bi-nucleated cells with the new centrin arm shorter than 1.0µm were significantly increased after RNAi induction (Fig. 4C). These results suggest that WDR2 is required to maintain centrin arm integrity. We next asked whether knockdown of WDR2 disrupted the integrity of the fishhook-like structure of the hook complex, and by immunofluorescence microscopy using the anti-TbMORN1 antibody, we showed that the old fishhook-like structure remained its typical morphology, but the new fishhook-like structure lost the shank part of the structure in ∼65% of the 2N2K cells and ∼86% of the 2N1K cells after RNAi (Fig. 4D, E), indicating the disruption of the morphology of the fishhook-like structure. Together, these results demonstrated that WDR2 is required for hook complex assembly by maintaining its morphology and integrity.

**Figure 4.**
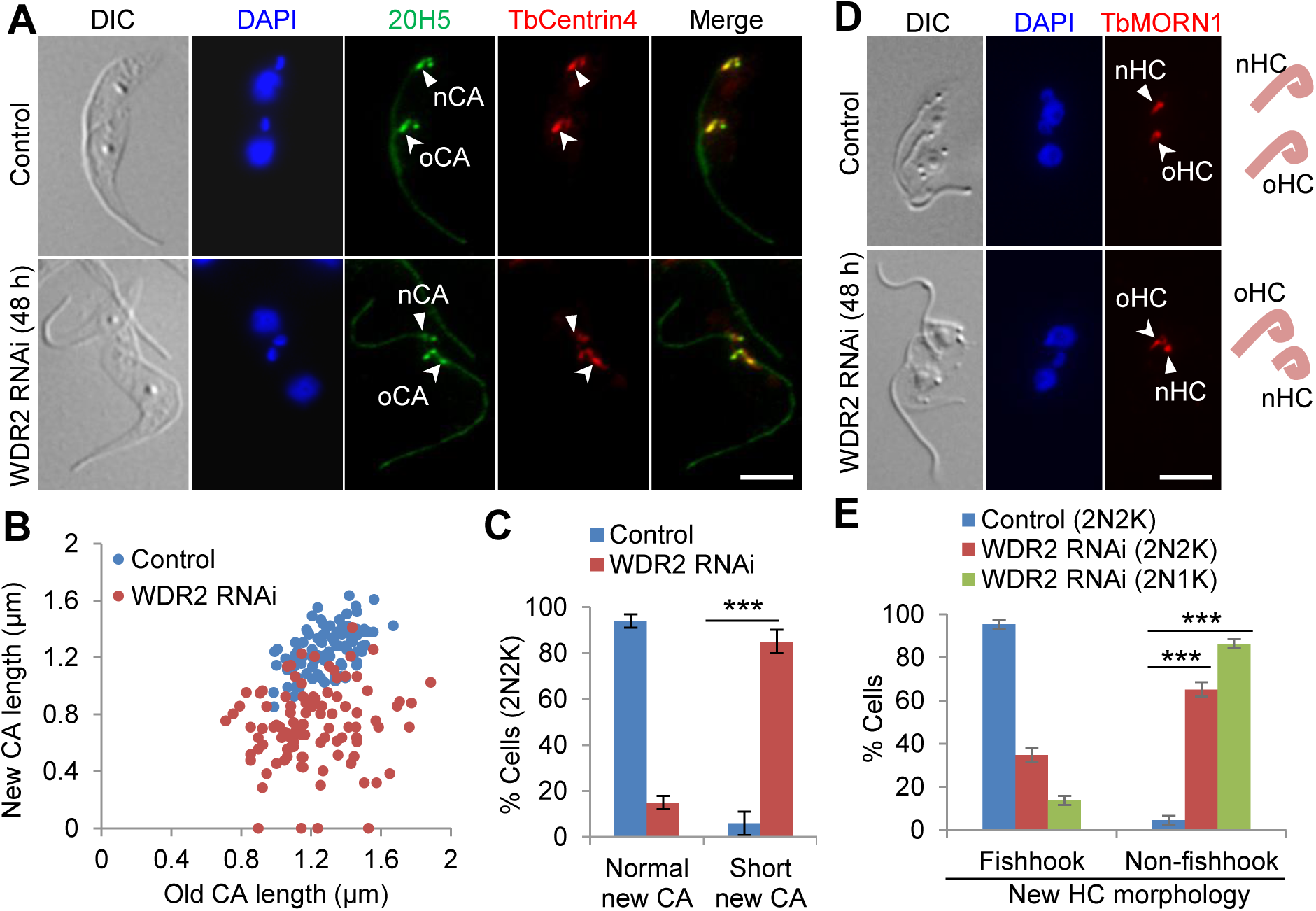
Knockdown of WDR2 impairs hook complex integrity. (**A**). Immunofluorescence microscopy to examine the centrin arm with the anti-TbCentrin4 antibody and the pan-centrin antibody 20H5. *Solid arrowheads* indicate the new centrin arm (nCA), and *open arrowheads* indicate the old centrin arm (oCA). Scale bar: 5 μm. **(B)**. Measurement of the length of the new centrin arm and the old centrin arm in control and WDR2 RNAi cells. **(C)**. Quantitation of bi-nucleated cells with normal-length centrin arm or short-length centrin arm in control and WDR2 RNAi cells. Error bars indicate S.D. from three independent experiments. ***, *p*<0.001. (**D**). Immunostaining of the hook complex with anti-TbMORN1 antibody. *Solid arrowheads* indicate the new hook complex (nHC), and *open arrowheads* indicate the old hook complex (oHC). Depicted on the right is the morphology of the new and the old hook complexes marked by anti-TbMORN1 antibody. Scale bar: 5 μm. (**E**). Quantitation of bi-nucleated cells with the hook complex displaying a typical fishhook-like morphology or a non-fishhook-like morphology in control and WDR2 RNAi cells. Error bars indicate S.D. from three independent experiments. ***, *p*<0.001.

### Knockdown of WDR2 impairs Golgi biogenesis

To investigate whether WDR2 RNAi also disrupts Golgi biogenesis, like its interacting partner KIN-G (Zhou et al., 2024), we performed co-immunofluorescence microscopy using the anti-TbGRASP antibody to detect the Golgi matrix protein TbGRASP and the anti-HA antibody to detect the 3HA-tagged CAAP1, a Golgi peripheral protein (Zhou et al., 2024), and then counted the number of Golgi in control and WDR2 RNAi cells (Fig. 5A, B). Knockdown of WDR2 resulted in a ∼50% reduction of the bi-nucleated cells with two, three, or four Golgi and a corresponding increase of bi-nucleated cells with more than four Golgi from ∼4% to ∼52% (Fig. 5B). Knockdown of WDR2 caused similar effects on the Sec13-marked ERES (ER exit site), a specialized ER zone involved in the ER-to-Golgi cargo transport (Kurokawa and Nakano, 2019), producing many bi-nucleated cells with more than four ERES foci (Fig. 5C). Many of the Golgi detected in the WDR2 RNAi cells were smaller in size, having weaker TbGRASP and CAAP1 signal (Fig. 5A), so were some of the ERESs, which had weaker Sec13 signal (Fig. 5C). Such smaller Golgi were also detected in control cells (Fig. 5A, arrows), as reported previously (de Graffenried et al., 2008; He et al., 2004), which often disappear upon cytokinesis, likely being integrated into the existing Golgi or being disassembled (He et al., 2004). Because the centrin arm determines the assembly site for new Golgi or regulates the size of the new Golgi (de Graffenried et al., 2008; He et al., 2005) and WDR2 disrupted centrin arm integrity (Fig. 4A-C), we wondered whether the shorter new centrin arm in WDR2 RNAi cells associates with a smaller Golgi. Thus, we performed co-immunofluorescence microscopy with the anti-TbGRASP antibody to label the Golgi and 20H5 to label the centrin arm (Fig. 5D). In the non-induced control cells, the old centrin arm-associated and the new centrin arm-associated Golgi had similar TbGRASP signal intensity, but in the WDR2 RNAi cells, the new centrin arm-associated Golgi had weaker TbGRASP signal than the old centrin arm-associated Golgi (Fig. 5D, E). Together, these results demonstrated that WDR2 depletion disrupted Golgi biogenesis, likely by impairing the maturation of the new centrin arm-associated Golgi. This phenotype is similar to the knockdown of KIN-G (Zhou et al., 2024) and the knockdown of TbPLK (de Graffenried et al., 2008), suggesting that WDR2, KIN-G, and TbPLK may function in the same pathway to regulate Golgi biogenesis.

**Figure 5.**
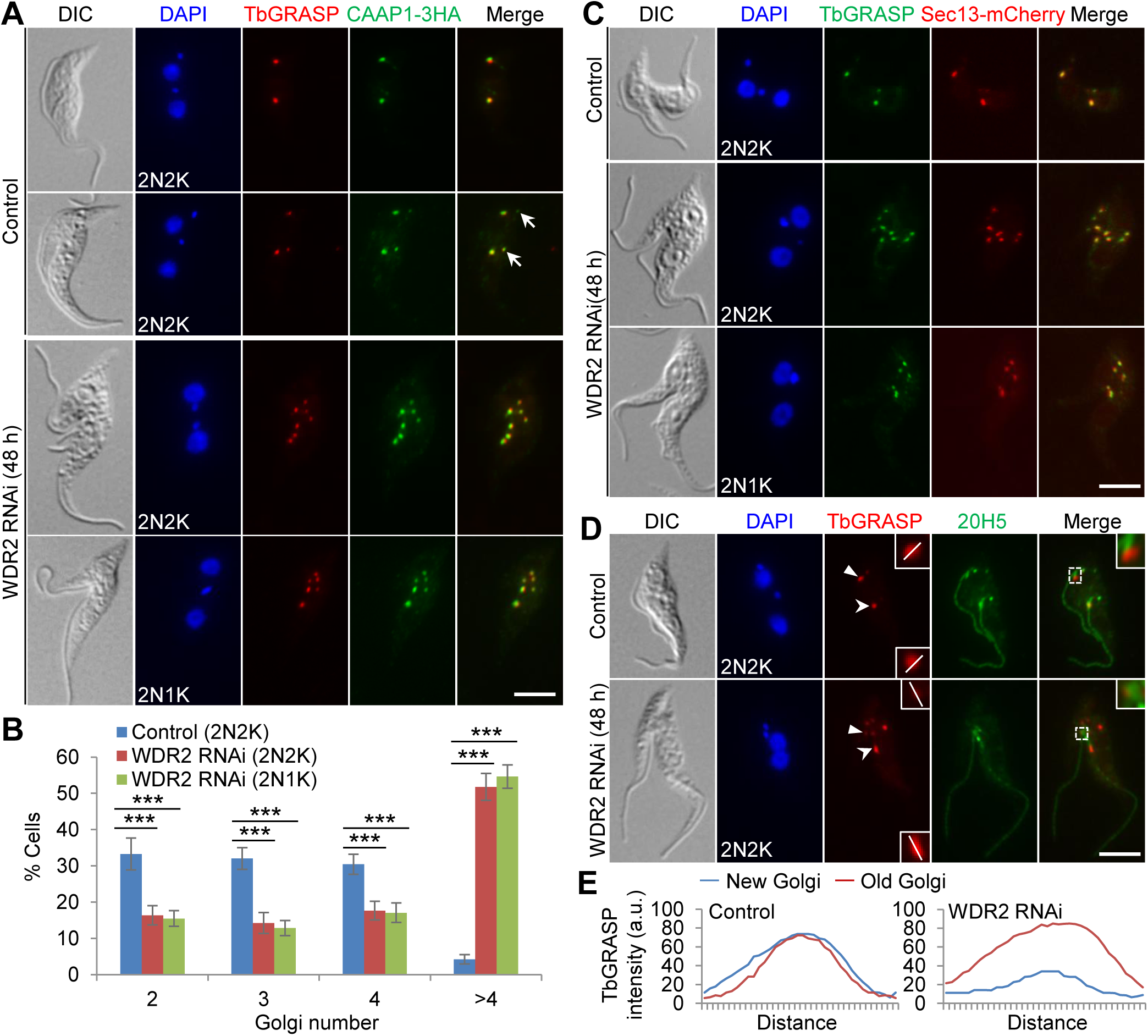
WDR2 is required for Golgi biogenesis. (**A**). Immunostaining of TbGRASP and CAAP1 in control and WDR2 RNAi cells. TbGRASP was detected by the anti-TbGRASP antibody, and endogenously 3HA-tagged CAAP1 was detected by the anti-HA antibody. *Arrows* indicate small Golgi. Scale bar: 5 μm. (**B**). Quantitation of the numbers of Golgi in control and WDR2 RNAi-induced bi-nucleated cells. Error bars indicate S.D. from three independent experiments. ***, *p*<0.001. (**C**). Fluorescence microscopic analysis of Sec13-mCherry in control and WDR2 RNAi cells. TbGRASP served as a marker for the Golgi. Scale bar: 5 μm. (**D**). Effect of WDR2 RNAi on Golgi-centrin arm association. Centrin arm was immunostained by the 20H5 antibody, and Golgi was detected by the anti-TbGRASP antibody. *Solid arrowheads* indicate the new centrin arm-associated Golgi, and *open arrowheads* indicate the old centrin arm-associated Golgi. The white line indicates the transect for quantitative measurement of TbGRASP signal shown in panel **E**. Insets in the merged panel show the zoom-in view of the selected region. Scale bar: 5 μm. (**E**). Quantitation of TbGRASP signal intensity in the centrin arm-associated new and old Golgi in control and WDR2 RNAi cells presented in panel **D**.

### Knockdown of WDR2 disrupts KIN-G localization and destabilizes KIN-G

Because WDR2 forms a complex with KIN-G (Fig. 1) and the phenotype of WDR2 knockdown mimicked that of KIN-G knockdown (Figs. 2-5), we asked whether WDR2 regulates KIN-G or vice versa. Immunofluorescence microscopy showed that knockdown of WDR2 disrupted KIN-G localization at the centrin arm (Fig. 6A). After WDR2 RNAi for 24 h, KIN-G signal at the new centrin arm either became weaker or was undetectable in ∼44% of the 1N2K cells and ∼45% of the 2N2K cells (Fig. 6A, B). Moreover, KIN-G was undetectable at both the new and the old centrin arms in ∼11% each of the 1N2K and 2N2K cells (Fig. 6A, B). After WDR2 RNAi for 48 h, KIN-G was undetectable at both the new and the old centrin arms in ∼83% of the 1N2K cells and ∼97% of the 2N2K cells (Fig. 6A, B). These results suggest that WDR2 is required for the recruitment of KIN-G to the new centrin arm and for the maintenance of KIN-G at the old centrin arm. Western blotting showed that the level of KIN-G protein was gradually decreased after WDR2 RNAi induction, and treatment of WDR2 RNAi-induced cells (day 2) with the proteasome inhibitor MG132 for 8 h restored KIN-G protein level (Fig. 6C). Immunofluorescence microscopy showed that KIN-G remained undetectable at both the new and the old centrin arms in the WDR2 RNAi cells (day 2) treated with MG132, albeit the overall cytosolic KIN-G signal was substantially increased (Fig. 6A), suggesting that KIN-G was degraded in the cytosol of WDR2 RNAi cells. Conversely, immunofluorescence microscopy showed that WDR2 was still detected at the centrin arm in KIN-G RNAi cells (Fig. 6D), and western blotting showed that WDR2 protein level was unaffected by KIN-G RNAi (Fig. 6E). Altogether, these results suggest that WDR2 is not a KIN-G cargo, but instead it functions to recruit and stabilize KIN-G.

**Figure 6.**
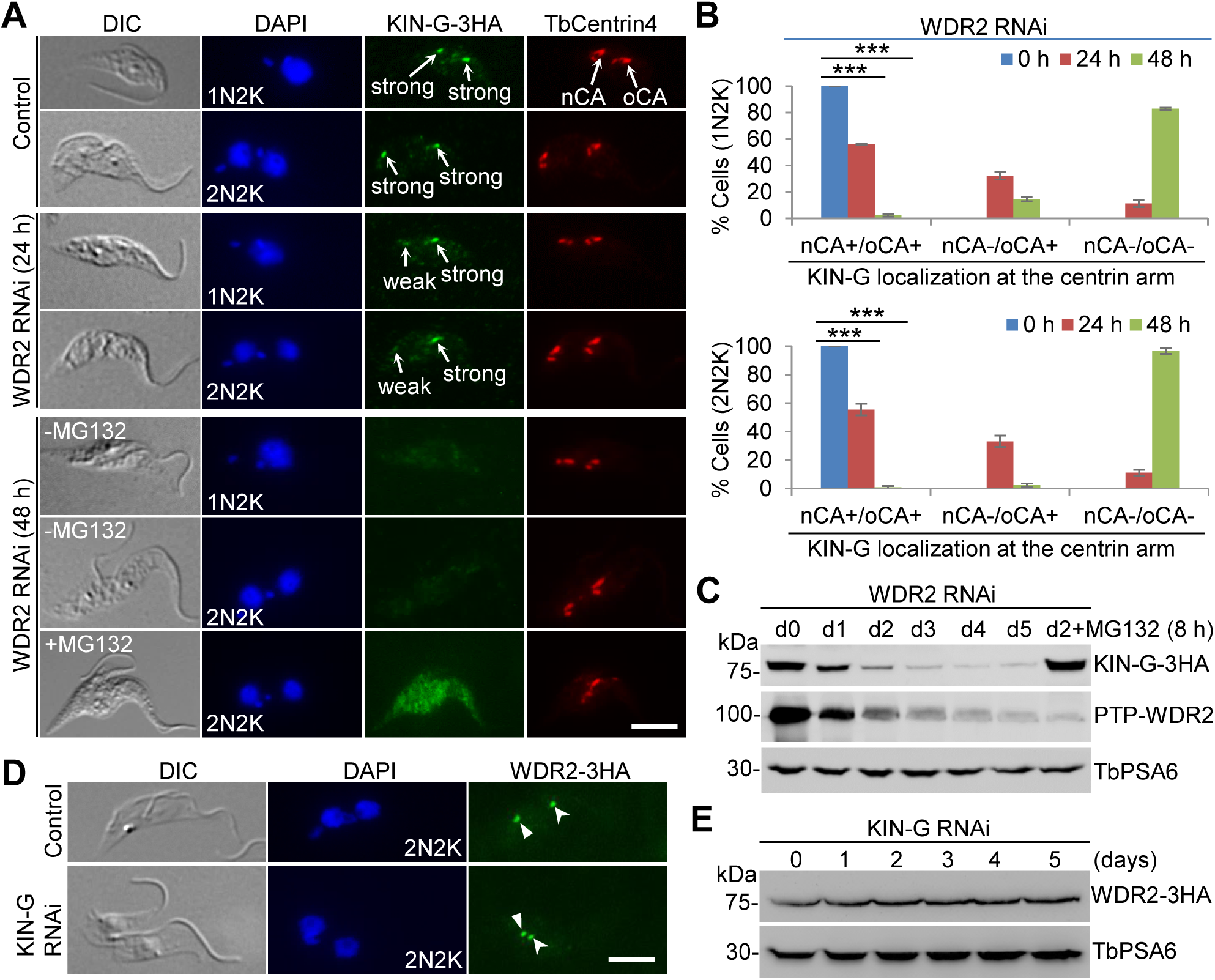
WDR2 is required for recruitment and maintenance of KIN-G at the centrin arm. (**A**). Immunofluorescence microscopic analysis of KIN-G-3HA in control, WDR2 RNAi cells, and WDR2 RNAi cells treated with MG132. nCA: new centrin arm; oCA: old centrin arm. Scale bar: 5 μm. (**B**). Quantitation of 1N2K and 2N2K cells with different localization patterns of KIN-G at the new and the old centrin arms in control and WDR2 RNAi cells. Error bars indicate S.D. from three independent experiments. nCA+/oCA+: KIN-G-positive at both the new and the old centrin arms; nCA-/oCA+: KIN-G-negative at the new centrin arm but positive at the old centrin arm; nCA-/oCA-: KIN-G-negative at both the new and the old centrin arms. ***, *p*<0.001. (**C**). Western blotting to detect the level of endogenously 3HA-tagged KIN-G before and after WDR2 RNAi induction and in WDR2 RNAi-induced cells (2 d) treated with MG132 (8 h). TbPSA6 served as a loading control. (**D**). Immunofluorescence microscopic analysis of WDR2-3HA in control and KIN-G RNAi cells. *Solid arrowhead* and *open arrowhead* indicate the WDR2 fluorescence signal at the new and the old centrin arms, respectively. Scale bar: 5 μm. (**E**). Western blotting to detect the level of endogenously 3HA-tagged WDR2 before and after KIN-G RNAi induction. TbPSA6 served as a loading control.

### Determination of WDR2 structural motifs required for interaction with KIN-G

The structural motifs in WDR2 were predicted by homology modeling and AlphaFold2 prediction (Fig. 7A-C). WDR2 contains in its C-terminal domain (CTD) a WD40 domain composed of seven WD40 repeats, which adopts a circularized, seven-bladed β-propeller structure, like a circular solenoid (Fig. 7A-C). WD40 domain is known to serve as a rigid scaffold for protein-protein interaction, and WD40 domain-containing proteins often function in coordinating multi-protein complex assembly (Smith et al., 1999). Thus, the WD40 domain in WDR2 may play roles in mediating protein complex assembly in trypanosomes. AlphaFold2 also predicted two domains of unknown function (DUF) in the N-terminal domain (NTD) of WDR2, each of which comprises two β-sheets and three α-helices (Fig. 7A-C). Homology modeling by SWISS-MODEL (Arnold et al., 2006; Biasini et al., 2014) detected a PB1 (Phox and Bem1) domain in the middle domain (MD) of WDR2, which comprises about 80 amino acids and exhibits a ubiquitin-like β-grasp fold with two α-helices and five β-sheets (Fig. 7A-C). The PB1 domain is evolutionarily conserved among animals, fungi, plants, and protozoa, and functions as a protein-protein interaction module through PB1-mediated heterodimerization/homodimerization or interaction with other protein domains (Sumimoto et al., 2007). Within the middle domain of WDR2, a short coiled-coil (CC) motif was detected between DUF2 and PB1 (Fig. 7A, B).

**Figure 7.**
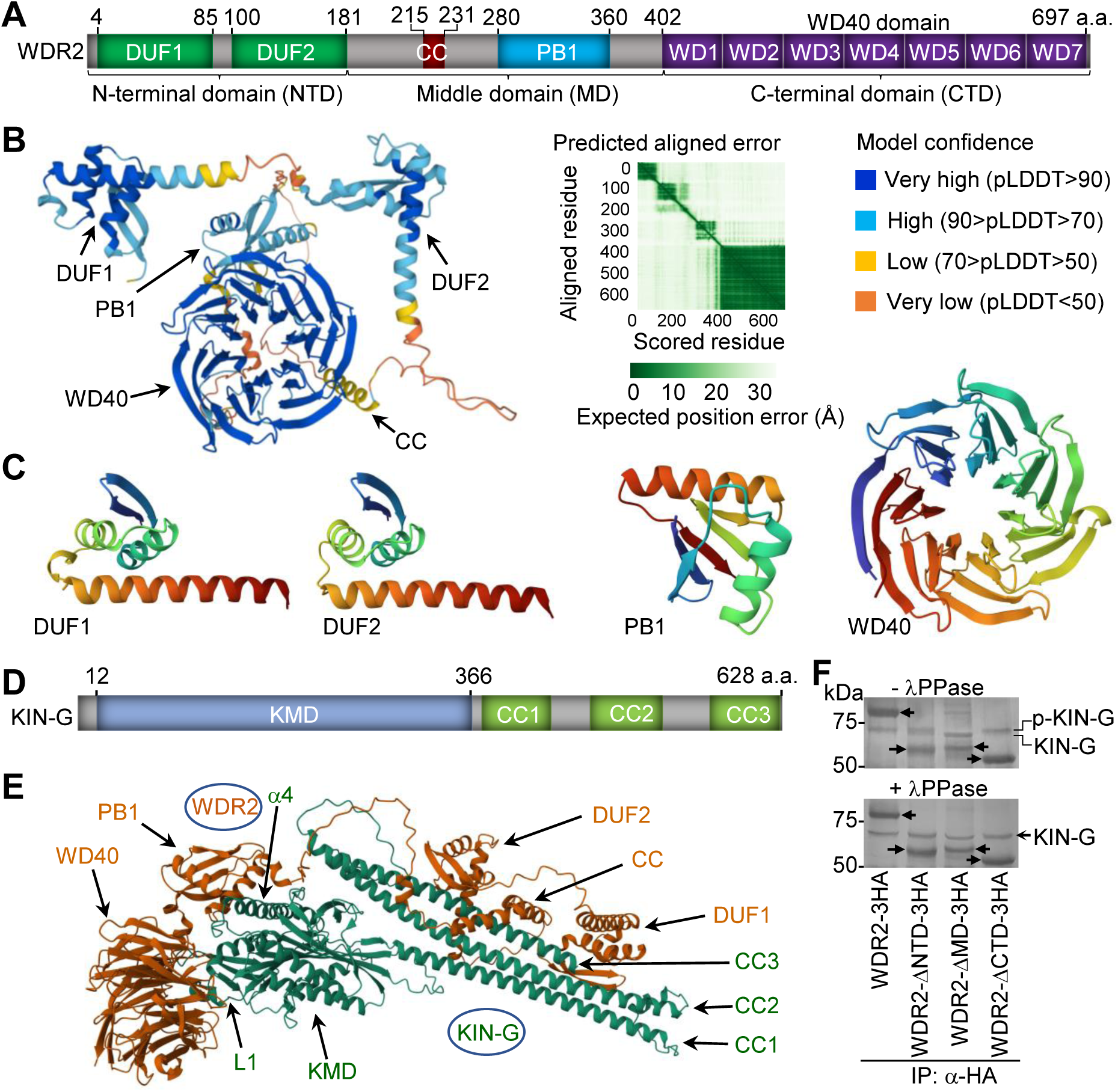
Structural domains of WDR2 required for interaction with KIN-G. (**A**). Schematic drawing of the structural domains in WDR2. DUF: domain of unknown function; CC: coiled coil; PB1: Phox and Bem1; WD: Tryptophan (W)-Aspartate D); NTD: N-terminal domain; MD: middle domain; CTD: C-terminal domain. (**B**). Structure of WDR2 predicted by AlphaFold2. The structural domains were indicated. (**C**). Structural domains predicted by SWISS-MODEL or AlphaFold2. The templates used for structural modeling of the PB1 domain and the WD40 domain in WDR2 are 2PPH and 8OKF, respectively. (**D**). Schematic drawing of the structural domains in KIN-G. KMD: kinesin motor domain; CC: coiled coil. (**E**). Structure of the WDR2-KIN-G complex predicted by AlphaFold3. The structural domains in each protein and the Loop #1 (L1) and the α-helices #4 (α4) of the KMD domain of KIN-G were indicated. (**F**). Co-immunoprecipitation to detect the *in vivo* interaction between native KIN-G and wild-type and the domain-deletion mutants of WDR2. Protein bands were stained by silver staining and identified by mass spectrometry. Solid arrows indicate the ectopically expressed, 3HA-tagged WDR2 and its domain-deletion mutants. p-KIN-G: phosphorylated KIN-G; λPPase: Lambda protein phosphatase.

As WDR2 forms a complex with KIN-G, which contains a kinesin motor domain (KMD) in the N-terminus and three CC motifs (CC1-CC3) in the C-terminus (Fig. 7D), we predicted the structure of the WDR2-KIN-G complex using AlphaFold3 (Abramson et al., 2024), which showed that the two proteins may interact via multiple domains (Fig. 7E). The WD40 domain of WDR2 may interact with the loop #1 (L1) of the KMD of KIN-G, the CC motif and the DUF2 domain of WDR2 may interact with the CC3 motif of KIN-G, and the PB1 domain of WDR2 may interact with the α-helix #4 (α4) of the KMD of KIN-G (Fig. 7E). To test the potential contribution of WDR2 structural domains to the interaction with KIN-G, we ectopically expressed wild-type WDR2 and three domain-deletion mutants, WDR2-ΔNTD, WDR2-ΔMD, and WDR2-ΔCTD, each of which was tagged with a C-terminal triple HA epitope, in the WDR2-3’UTR RNAi cell line (see below for details), and then performed co-immunoprecipitation followed by mass spectrometry. Wild-type WDR2, WDR2-ΔNTD, and WDR2-ΔCTD each pulled down a protein of ∼70 kDa (Fig. 7F), which was identified as KIN-G by mass spectrometry (Supplemental Fig. S1 and Supplemental Table S2). However, WDR2-ΔMD pulled down a protein with a slightly smaller molecular mass than that pulled down by wild-type WDR2 and the other two WDR2 mutants (Fig. 7F), and this protein was also identified as KIN-G by mass spectrometry (Supplemental Fig. S1 and Supplemental Table S3), suggestive of a potential non-phosphorylated form of KIN-G pulled down by WDR2-ΔMD only. To test this possibility, trypanosome cell lysate was treated with Lambda protein phosphatase (λPPase) prior to co-immunoprecipitation, and wild-type WDR2 and all the three WDR2 mutants pulled down the KIN-G protein of the same molecular mass (Fig. 7F). These results suggest two possibilities. First, KIN-G is de-phosphorylated in the WDR2-ΔMD mutant cells, but the dephosphorylated KIN-G still interacts with WDR2-ΔMD. Second, WDR2-ΔMD only interacts with de-phosphorylated KIN-G, whereas wild-type WDR2 and the other two WDR2 mutants interact with phosphorylated KIN-G. Nonetheless, these results suggest that none of the three domains in WDR2 are exclusively required for the interaction with KIN-G and all of the three domains (NTD, MD, and CTD) mediate WDR2-KIN-G interaction, in agreement with the interaction interfaces predicted by AlphaFold3 (Fig. 7E).

### Structural motifs required for WDR2 cellular function and KIN-G localization

To investigate the potential function of the DUF domains in the NTD, the CC and the PB1 in the MD, and the WD40 domain in the CTD, we deleted the NTD, the MD, or the CTD of WDR2 (Fig. 8A) for complementation of WDR2 deficiency. To this end, we first generated a WDR2-3’UTR RNAi cell line by targeting against the 3’UTR of WDR2, and western blotting demonstrated the knockdown of endogenous, PTP-tagged WDR2 (Fig. 8B). This knockdown caused growth defects (Fig. 8C), similar to the RNAi against the coding region of WDR2 (Fig. 2B). Subsequently, we ectopically expressed wild-type and the domain-deletion mutants of WDR2, each of which was tagged with a C-terminal triple HA epitope, in the WDR2-3’UTR RNAi cell line (Fig. 8D). Expression of wild-type WDR2 was able to restore the growth defects of WDR2-3’UTR RNAi cells, whereas expression of WDR2-ΔNTD partially restored the growth defects, and expression of WDR2-ΔMD or WDR2-ΔCTD failed to restore the growth defects of the WDR2-3’UTR RNAi cells (Fig. 8E). These results suggest that the NTD is important for WDR2 function, whereas the MD and the CTD are essential for WDR2 function.

**Figure 8.**
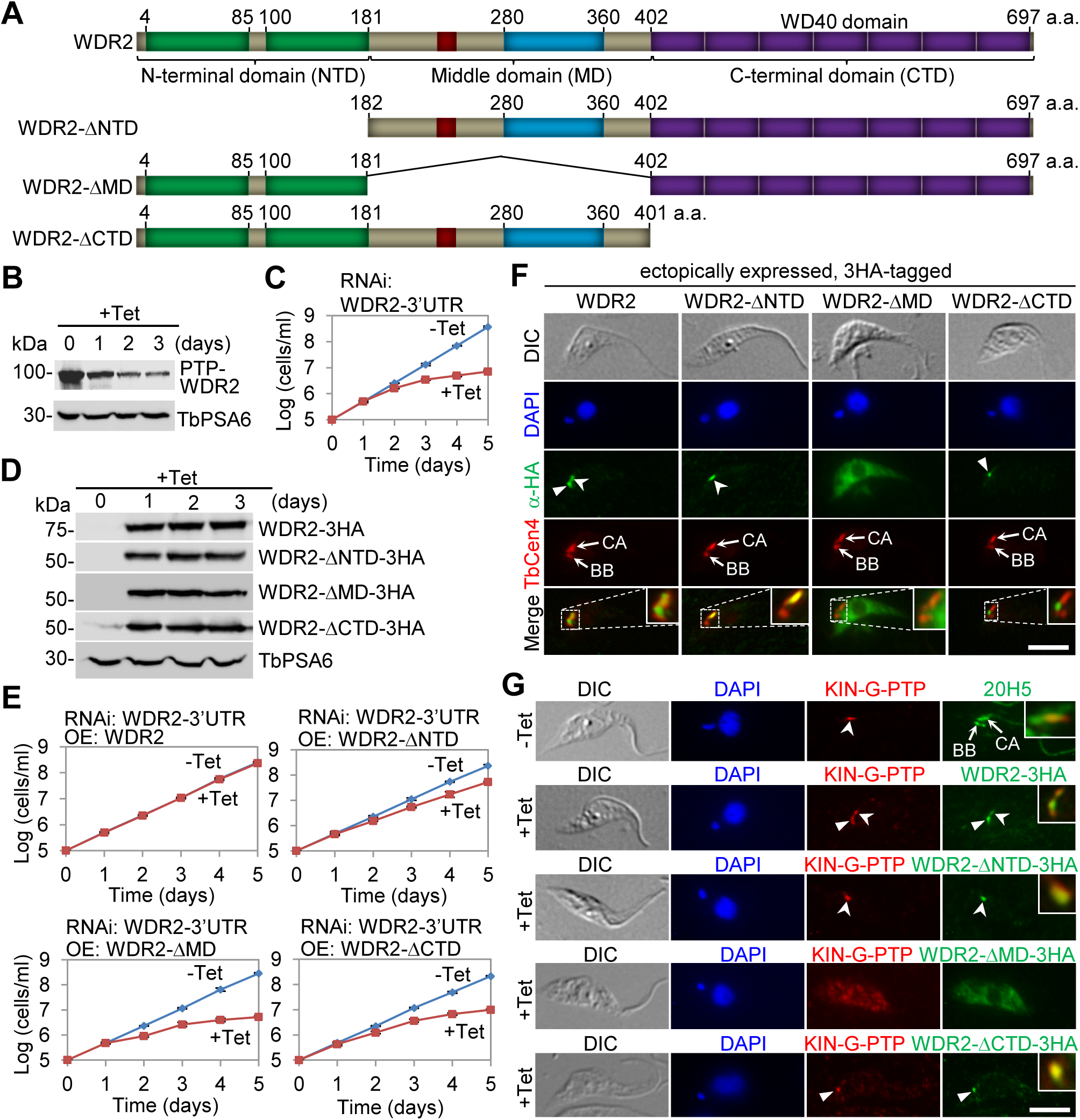
Structural domains of WDR2 required for WDR2 cellular function. (**A**). Schematic drawing of WDR2 and its domain-deletion mutants used for genetic complementation. (**B**). Western blotting to detect endogenously PTP-tagged WDR2 in WDR2-3’UTR RNAi cell line induced for 3 days. TbPSA6 served as a loading control. (**C**). Growth curve of the WDR2-3’UTR RNAi cell line. (**D**). Western blotting to detect ectopically expressed, 3HA-tagged wild-type and mutant WDR2 in the WDR2-3’UTR RNAi cell line induced for 3 days. TbPSA6 served as a loading control. (**E**). Growth curves of WDR2-3’UTR RNAi complementation cell lines expressing wild-type or the domain-deletion mutants of WDR2. (**F**). Subcellular localization of ectopically expressed, 3HA-tagged WDR2, WDR2-ΔNTD, WDR2-ΔMD, and WDR2-ΔCTD. Cells were co-immunostained with the anti-HA antibody and the anti-TbCentrin4 antibody. BB: basal body; CA: centrin arm. *Solid arrowheads* indicate the WDR2 signal at the MtQ proximal end, and *open arrowheads* indicate the WDR2 signal at the centrin arm. Insets show the zoom-in view of the selected region. Scale bar: 5 μm. (**G**). Co-immunostaining of endogenously expressed, PTP-tagged KIN-G and ectopically expressed, 3HA-tagged wild-type and the domain-deletion mutants of WDR2. *Solid arrowheads* indicate the WDR2 signal at the MtQ proximal end, and *open arrowheads* indicate the WDR2 signal at the centrin arm. Insets show the zoom-in view of the co-localization. Scale bar: 5 μm.

We next examined the subcellular localization of wild-type and the three domain-deletion mutants of WDR2 by immunofluorescence microscopy. Ectopically expressed, 3HA-tagged WDR2 localized to the centrin arm but extended to the region between the basal body and the centrin arm, reminiscent of the proximal end of the MtQ (Fig. 8F), in contrast to the endogenously 3HA-tagged WDR2, which was detected only at the centrin arm (Fig. 1E). Similarly, ectopically expressed, 3HA-tagged KIN-G was detected at both the centrin arm and the MtQ proximal end (Zhou et al., 2024), in contrast to the endogenously 3HA-tagged KIN-G, which was detected only at the centrin arm (Figs. 1E and 6A). The discrepancy could be due to the ectopic expression that produced more proteins to be spread to the MtQ proximal end. Nonetheless, the WDR2-ΔNTD mutant was only detected at the centrin arm, the WDR2-ΔCTD mutant was only detected at the MtQ proximal end, and the WDR2-ΔMD mutant was mis-localized to the cytosol (Fig. 8F). These results suggest that the CTD is required for localization of the ectopically overexpressed WDR2 to the centrin arm, the NTD is required for localization to the MtQ proximal end, and the MD is required for localization to both the centrin arm and the MtQ proximal end.

Because WDR2 knockdown disrupted KIN-G localization (Fig. 6), we wondered whether WDR2 can actively target KIN-G to its subcellular location. To test this possibility, we examined the localization of endogenous, PTP-tagged KIN-G in WDR2-3’UTR RNAi cells expressing ectopic WDR2 and its domain-deletion mutants, which localized to both the centrin arm and the MtQ proximal end, the centrin arm, the cytosol, or the MtQ proximal end, respectively (Fig. 8F). In non-induced cells, endogenously PTP-tagged KIN-G was detectable only at the centrin arm (Fig. 8G), as expected. However, in cells induced to ectopically express WDR2, KIN-G co-localized with WDR2 at both the centrin arm and the MtQ proximal end (Fig. 8G). In cells induced to ectopically express WDR2-ΔNTD, KIN-G co-localized with WDR2-ΔNTD at the centrin arm (Fig. 8G). In cells induced to ectopically express WDR2-ΔMD, KIN-G was mis-localized to cytosol (Fig. 8G), and in cells induced to ectopically express WDR2-ΔCTD, KIN-G co-localized with WDR2-ΔCTD at the MtQ proximal end (Fig. 8G). Thus, the endogenously PTP-tagged KIN-G was targeted to the same locations where ectopically expressed WDR2 and its domain-deletion mutants were localized, suggesting that WDR2 can actively recruit KIN-G.

## Discussion

The centrin arm structure in *T. brucei* is defined by two centrin proteins, TbCentrin2 and TbCentrin4, and was originally termed the bilobe structure involved in Golgi biogenesis, with one lobe associating with the existing Golgi and the other associating with the growing Golgi (He et al., 2005). Based on the findings from knockdown of TbCentrin2, the centrin arm is hypothesized to determine the site where the new Golgi is assembled (He et al., 2005). However, knockdown of TbCentrin4 had no effect on centrin arm assembly and Golgi biogenesis, but it disrupted the coordination of karyokinesis and cytokinesis (Shi et al., 2008). Three other proteins, TbPLK, WDR1, and KIN-G, also localize to the centrin arm. TbPLK regulates centrin arm assembly and Golgi biogenesis, but its regulation of Golgi biogenesis is opposite to that of TbCentrin2, despite that TbCentrin2 is a substrate of TbPLK (de Graffenried et al., 2008). WDR1 is required for centrin arm assembly and basal body segregation, and it plays these roles by controlling the abundance of TbPLK at the centrin arm and the basal body (Hu et al., 2017). Whether WDR1 regulates Golgi biogenesis was not investigated previously, but given its role in regulating TbPLK, it is likely that WDR1 also regulates Golgi biogenesis through an indirect means via TbPLK. KIN-G is an orphan kinesin, and it promotes the biogenesis of the hook complex and the Golgi (Zhou et al., 2024), similar to TbPLK (de Graffenried et al., 2008). KIN-G plays a role in Golgi biogenesis by recruiting the Golgi peripheral protein CAAP1, which promotes the association of Golgi with the centrin arm (Zhou et al., 2024). As a microtubule plus end-directed motor protein, KIN-G may transport cargos along the MtQ to their destination at the centrin arm and the Golgi to facilitate hook complex and Golgi biogenesis. All in all, it appears that the above-mentioned centrin arm-localized proteins play diverse functions, although some of them may act in the same pathway.

We have attempted to identify the potential cargo proteins of KIN-G by proximity biotinylation and have identified a subset of KIN-G-proximal proteins, including three centrin arm-localized proteins (TbCentrin2, TbCentrin4, and WDR2), four hook complex-localized proteins (Hypo.1-Hypo.4), the hook shank-localized protein TbSmee1, and two MtQ proximal end-localized proteins (Hypo.8 and Hypo.10) (Fig. 1B, C). Although the hook complex-localized proteins and MtQ proximal end-localized proteins are in close proximity to KIN-G, they are unlikely to be cargo proteins of KIN-G due to the enrichment of KIN-G in the distal part of the centrin arm (Zhou et al., 2024). We postulate that the most likely cargo proteins of KIN-G are those proteins that localize to the centrin arm-Golgi region, including the hook shank-localized TbSmee1, Golgi peripheral protein CAAP1, and centrin arm-localized proteins. The centrin arm-localized WDR2 was not affected by KIN-G RNAi (Fig. 6), ruling out its candidacy as a KIN-G cargo protein. TbSmee1 was only slightly reduced in some (∼20%) of the KIN-G RNAi cells (Zhou et al., 2024); thus, it is unlikely to be a cargo for KIN-G. In contrast, CAAP1 and TbCentrin4 were significantly reduced from the new Golgi and the new centrin arm, respectively, in KIN-G knockdown cells (Zhou et al., 2024), suggesting that they are potential cargo proteins of KIN-G, although their candidacy as KIN-G cargos remains to be verified by additional experimental approaches. Whether KIN-G transports any other types of cargos, such as vesicles, lipids, and/or organelles, remains to be explored, and is beyond the scope of this study.

We have provided evidence to support a regulatory role for WDR2 in the recruitment and maintenance of KIN-G at the centrin arm (Fig. 6). Knockdown of WDR2 disrupted the localization of KIN-G to both the new and the old centrin arms, resulting in the eventual degradation of KIN-G in the cytosol (Fig. 6). Since the new centrin arm undergoes *de novo* biogenesis during early cell cycle, the disrupted localization of KIN-G to the new centrin arm by WDR2 knockdown (Fig. 6A, B) suggests that WDR2 is required for recruiting KIN-G to the new centrin arm. This notion was further supported by the finding that native KIN-G interacted with and co-localized with the ectopically overexpressed wild-type or the domain-deletion mutants of WDR2 (Figs. 7F and 8G), which demonstrated an active action of WDR2 in recruiting KIN-G. Further, since KIN-G had already been recruited to the old centrin arm prior to WDR2 RNAi induction, the disrupted localization of KIN-G at the old centrin arm (Fig. 6A, B) suggests that WDR2 is also required for maintaining KIN-G at the old centrin arm. It should be noted that the observed loss of KIN-G signal at the old centrin arm after WDR2 RNAi can also be due to defects in recruitment. In this scenario, the 2N2K cells that failed to recruit KIN-G to the new centrin arm divided to produce a 1N1K new-flagellum daughter cell that further progressed through the cell cycle to become a 1N2K cell and then a 2N2K cell, in which KIN-G was lost at both the new and the old centrin arms. Because this 1N1K new-flagellum daughter cell has a long unattached flagellum, its derived 1N2K cell and 2N2K cell will have long unattached new and old flagella. However, we examined the 1N2K and 2N2K cells that had a normal old flagellum and found that they lost KIN-G signal at the old centrin arm (Fig. 6A). This result suggests that WDR2 plays a role in maintaining KIN-G at the old centrin arm. Together, these results uncovered a mechanistic role for WDR2 in recruiting and maintaining KIN-G at the centrin arm. Because KIN-G was degraded in the cytosol of the WDR2 RNAi cells (Fig. 6A, C), whereas the cytosolically mis-localized KIN-G in the WDR2-3’UTR RNAi cells expressing WDR2-ΔMD was stable (Figs. 7F and 8G), it suggests that interaction with WDR2 stabilizes KIN-G. Knockdown of WDR2 phenocopies all of the defects caused by knockdown of KIN-G in procyclic trypanosomes. First, depletion of WDR2 arrested cytokinesis progression, producing a normal-sized daughter cell and a small-sized daughter cell with a long unattached flagellum, due to the inhibition of new FAZ elongation (Fig. 2). Secondly, knockdown of WDR2 disrupted the positioning of the newly assembled flagellum and the segregation of flagellum-associated cytoskeletal structures (Fig. 3). Thirdly, deficiency in WDR2 disrupted the typical morphology of the hook complex by reducing the length of the centrin arm and eliminating the shank part of the fishhook-like structure (Fig. 4). Finally, knockdown of WDR2 impaired Golgi biogenesis by disrupting the maturation of the new centrin arm-associated Golgi (Fig. 5). Since WDR2 localizes to the centrin arm, the primary defect caused by WDR2 knockdown is the disruption of hook complex integrity, which led to secondary defects on the elongation of the new FAZ and the positioning of the new flagellum. Consequently, the defects in FAZ elongation and flagellum positioning caused the mis-positioning of the cell division plane, leading to asymmetrical cytokinesis (Fig. 2). All of these cellular defects caused by WDR2 RNAi could be attributed to the disrupted localization and destabilization of KIN-G (Fig. 6). Thus, the primary function for WDR2 is to recruit KIN-G to the centrin arm for the latter to play its role in regulating hook complex and Golgi biogenesis.

The endogenous, epitope-tagged WDR2 and KIN-G were detected only at the centrin arm (Fig. 1E). However, the ectopically expressed, epitope-tagged WDR2 and KIN-G were detected at both the centrin arm and the MtQ proximal end (Fig. 8 and (Zhou et al., 2024)), similar to the endogenous, mNeoGreen-tagged WDR2 and KIN-G reported in the TrypTag project (Billington et al., 2023). It is likely that native WDR2 and KIN-G proteins localize to both the centrin arm and the MtQ proximal end, but for unknown reasons, the anti-HA and the anti-Protein A antibodies used in our work were not able to detect the two proteins at the MtQ proximal end. Nonetheless, localization of KIN-G to the centrin arm depends on its motor activity (Zhou et al., 2024), whereas localization of WDR2 to the centrin arm requires its WD40 domain (Fig. 8F). Intriguingly, localization of WDR2 to the MtQ proximal end requires the N-terminal DUF domains, and localization of WDR2 to both the centrin arm and the MtQ proximal end requires the CC and the PB1 motif in the middle domain (Fig. 8F). It is possible that WDR2 is first recruited to the MtQ proximal end and the centrin arm, where WDR2 recruits KIN-G. Alternatively, WDR2 and KIN-G may form a complex in the cytosol, and the complex is then recruited to the two structures in a WDR2-dependent manner.

The finding that wild-type WDR2 and the WDR2-ΔNTD and WDR2-ΔCTD mutants, but not the WDR2-ΔMD mutant, pulled down phosphorylated KIN-G from trypanosome cell lysate (Fig. 7F) suggests that either KIN-G is phosphorylated at the centrin arm or phosphorylation of KIN-G targets it to the centrin arm. In the cytosol, KIN-G is non-phosphorylated, but it is still capable of forming a complex with the WDR2-ΔMD mutant (Fig. 7F), suggesting that complex formation is independent of KIN-G phosphorylation. Because KIN-G localization depends on WDR2 (Figs. 6 and 8), it suggests that phosphorylation of KIN-G is unlikely to play a role in targeting itself to the centrin arm. The best candidate protein kinase responsible for KIN-G phosphorylation is TbPLK, which localizes to the centrin arm and is required for hook complex and Golgi biogenesis (de Graffenried et al., 2008), similar to the role of KIN-G. Whether TbPLK phosphorylates KIN-G and how the phosphorylation of KIN-G may impact the biochemical function of KIN-G, i.e., the motor activity and/or microtubule-binding activity, and the physiological function of KIN-G in regulating the biogenesis of the hook complex and the Golgi are currently under investigation.

In closing, we identified WDR2 as a KIN-G-interacting partner protein, which co-localizes with KIN-G at the centrin arm in the procyclic form of *T. brucei* and regulates KIN-G by recruiting the latter to the centrin arm, thereby promoting the biogenesis of the hook complex and the Golgi apparatus. The disruption of hook complex biogenesis further impaired FAZ elongation and flagellum positioning, leading to mis-placement of the cell division plane and, consequently, asymmetrical cytokinesis.

## Materials and Methods

### Structural modeling and AlphaFold prediction of protein structure

Structural modeling of WDR2 structural motifs was carried out by SWISS-MODEL (https://swissmodel.expasy.org/) (Arnold et al., 2006; Biasini et al., 2014). The WDR2 protein structure was obtained directly from AlphaFold protein structure database (https://alphafold.ebi.ac.uk/), and the two N-terminal domains of unknown function in WDR2 were predicted by AlphaFold2 (Jumper et al., 2021; Varadi et al., 2022). The structure of the WDR2-KIN-G complex was predicted by AlphaFold3 (Abramson et al., 2024) using the following webserver: https://alphafoldserver.com/.

### Trypanosome cell culture and RNAi

The procyclic *T. brucei* Lister427 strain was grown in SDM-79 medium containing 10% heat-inactivated fetal bovine serum (Millipore-Sigma) at 27°C, and the procyclic *T. brucei* strain 29-13 (Wirtz et al., 1999) was cultured in SDM-79 medium supplemented with 10% heat-inactivated fetal bovine serum, 15 µg/ml G418, and 50 µg/ml hygromycin at 27°C.

To generate the WDR2 RNAi cell line, a 453-bp DNA fragment (nucleotides 1407-1859) of the WDR2 gene was amplified from trypanosome genomic DNA by PCR and cloned into the pZJM vector (Wang et al., 2000). The resulting plasmid, pZJM-WDR2, was used to transfect the 29-13 strain by electroporation. Transfectants were selected with 2.5 µg/ml phleomycin, and successful transfectants were cloned by limiting dilution in a 96-well plate containing the SDM-79 medium supplemented with 20% fetal bovine serum and appropriate antibiotics. The KIN-G RNAi cell line was generated previously (Zhou et al., 2024).

RNAi was induced by incubating the RNAi cell line with 1.0 µg/ml tetracycline. Three clonal RNAi cell lines were chosen for phenotypic analysis, and because the three clonal cell lines showed almost identical phenotypes, only one clonal RNAi cell line was used for characterization throughout the work.

### *In situ* epitope tagging of proteins

Epitope tagging of WDR2, CAAP1, KIN-G, and Sec13 from their respective endogenous locus was carried out by the PCR-based epitope-tagging method reported previously (Shen et al., 2001). For WDR2 and KIN-G co-localization experiments, WDR2 was endogenously tagged with a C-terminal PTP epitope, and KIN-G was endogenously tagged with a C-terminal triple HA epitope in the same cell line. Transfectants were selected with appropriate antibiotics, and clonal cell lines were obtained by limiting dilution as described above.

### Generation of WDR2 RNAi complementation cell lines

To generate WDR2 RNAi complementation cell lines, we first created a WDR2-3’UTR RNAi cell line for ectopic expression of wild-type and the structure domain-deletion mutants of WDR2. To this end, a 564-bp fragment of the 3’UTR of WDR2 was cloned into the pZJM-PAC vector, and the resulting plasmid was electroporated into the procyclic 29-13 strain. Transfectants were selected with 1.0 µg/ml puromycin in addition to 15 µg/ml G418 and 50 µg/ml hygromycin, and clonal cell lines were generated by limiting dilution in a 96-well plate as described above. Subsequently, the full-length *WDR2* gene and the mutant *WDR2* gene lacking the sequence encoding the NTD, the MD, or the CTD were each cloned into the pLew100-3HA-BLE vector (Wei et al., 2014). The resulting plasmids, pLew100-WDR2-3HA-BLE, pLew100-WDR2-ΔNTD-3HA-BLE, pLew100-WDR2-ΔMD-3HA-BLE, and pLew100-WDR2-ΔCTD-3HA-BLE, were used to transfect the WDR2-3’UTR RNAi cell line. Transfectants were selected with 2.5 µg/ml phleomycin in additional to1.0 µg/ml puromycin, 15 µg/ml G418, and 50 µg/ml hygromycin, and then cloned by limiting dilution in a 96-well plate as described above. To induce WDR2-3’UTR RNAi and ectopic expression of WDR2-3HA and its domain-deletion mutants, cells were incubated with 1.0 μg/ml tetracycline.

### Co-immunoprecipitation, λPPase treatment, silver staining, and mass spectrometry

Trypanosomes cells expressing KIN-G-3HA, WDR2-PTP, or both KIN-G-3HA and WDR2-PTP at their respective endogenous locus were lysed in 0.5 ml immunoprecipitation (IP) buffer (25 mM Tris-HCl, pH 7.4, 100 mM NaCl, 1 mM DTT, 0.1% NP-40, 5% glycerol, and protease inhibitor cocktail) on ice for 30 min. Cell lysate was cleared by centrifugation at the highest speed in a table-top microcentrifuge. The cleared cell lysate was incubated with 25 μl settled IgG Sepharose beads (GE Healthcare) at 4°C for 60 min. After centrifugation at 4°C for 10 min in a microcentrifuge, the IgG Sepharose beads were washed five times with the IP buffer, and bound proteins were eluted by boiling the beads in 1x SDS sampling buffer for 5 min. Eluted proteins were separated by SDS-PAGE, transferred onto a PVDF membrane, and immunoblotted with anti-HA monoclonal antibody (clone HA-7, H9658, Sigma-Aldrich, 1:5,000 dilution) to detect 3HA-tagged KIN-G, and anti-Protein A polyclonal antibody (anti-ProtA; P3775, Sigma-Aldrich, 1:5,000 dilution) to detect PTP-tagged WDR2.

Trypanosome cells expressing ectopic WDR2-3HA or its domain-deletion mutants in the WDR2-3’UTR RNAi cell line were lysed in the IP buffer on ice for 30 min. For Lambda protein phosphatase (λPPase) treatment, 100 units of λPPase (New England Biolabs, Cat#: P0753), 1× NEBuffer for Protein MetalloPhosphatases, and 1 mM MnCl2 were added into the cell lysate and incubated at 30°C for 30 min. Cell lysate was cleared by centrifugation, the cleared supernatant was incubated with 15 μl settled anti-HA agarose beads (Millipore-Sigma) at 4°C for 60 min. Beads were then washed five times with the IP buffer, and bound proteins were eluted for SDS-PAGE followed by silver staining using the Pierce™ Silver Stain Kit (Cat# 24612, ThermoFisher Scientific) according to manufacturer’s instructions.

Co-immunoprecipitated protein bands were excised from the SDS-PAGE gel and analyzed by mass spectrometry using the USCF in-gel digestion protocol described in our previous publication (An et al., 2018). After overnight trypsin digestion, peptides were extracted from the gel with 50% acetonitrile and 5% formic acid, dried using speed-vac, and then reconstituted in 2% acetonitrile with 0.1% formic acid. Peptides were injected on to Thermo LTQ Orbitrap XL (Thermo-Fisher Scientific, Bremen, Germany) interfaced with an Eksigent nano-LC 2D plus ChipLC system (Eksigent Technologies, Dublin, CA). Raw data files were processed and searched using the Mascot search engine against the *T. brucei* proteome database. Mass spectrometry and data analysis were carried out at the Clinical and Translational Proteomics Service Center of the University of Texas Health Science Center at Houston.

### Proximity-dependent biotin identification and mass spectrometry

To identify proximal proteins of KIN-G and WDR2 by BioID, the full-length sequence of *KIN-G* and *WDR2* genes was each cloned into the pLew100-BirA*-HA vector (Hu et al., 2015b), and the resulting plasmid was linearized by NotI restriction digestion and used to transfect the procyclic 29-13 strain. Transfectants were selected with 2.5 µg/ml phleomycin and cloned by limiting dilution as described above. To express KIN-G-BirA*-HA or WDR2-BirA*-3HA, cells were incubated with 0.5 µg/ml tetracycline for 24 h and subsequently 50 µM biotin were added and further incubated for 24 h. Cells were harvested and treated with PEME buffer (100 mM PIPES pH 6.9, 2 mM EGTA, 1 mM MgSO4, 0.1 mM EDTA) containing 0.5% Nonidet P-40, and the cytosolic (soluble) and cytoskeletal (pellet) fractions were prepared by centrifugation. To solubilize cytoskeletal proteins, the cytoskeletal fraction was extracted with lysis buffer (0.4% SDS, 500 mM NaCl, 5 mM EDTA, 1 mM DTT, 50 mM Tris-HCl, pH 7.4), and the cytoskeletal extract was incubated with 500 µl pre-washed streptavidin-coated Dynabeads (Invitrogen) at 4°C for 4 h. The Dynabeads were washed five times with PBS and then five time with 50 mM ammonium bicarbonate before re-suspending in 100 mM ammonium bicarbonate. Finally, 10% DTT, 50% iodoacetamide, and 5% DTT were sequentially added to the Dynabeads re-suspension. The Dynabeads resuspension was treated with trypsin at 37°C for 16 hours, and trypsin-digested peptides were desalted and analyzed on an LTQ Orbitrap XL mass spectrometer (Thermo-Fisher Scientific) interfaced with an Eksgent nano-LC 2D plus chipLC system (Eksigent Technologies). Raw mass spectrometry data was searched against the *T. brucei* proteome database using the Mascot search engine. The search conditions used peptide tolerance of 10 ppm and MS/MS tolerance of 0.8 Da with the enzyme trypsin and two missed cleavages.

### Immunofluorescence microscopy

Trypanosome cells were settled onto glass coverslips, fixed with methanol at −20°C for 30 min, and then rehydrated with PBS for 10 min at room temperature. Cells on the coverslips were blocked with 3% BSA in PBS at room temperature for 1 h, and incubated at room temperature for 1 h with the following primary antibodies: anti-CC2D polyclonal antibody (1:1,000 dilution) (Zhou et al., 2011), L8C4 (anti-PFR2) monoclonal antibody (1:50 dilution) (Kohl et al., 1999), 20H5 monoclonal antibody (1:400 dilution) (He et al., 2005), anti-TbCentrin4/LdCentrin1 polyclonal antibody (1:1000 dilution), anti-TbMORN1 antibody (1:5000 dilution) (Morriswood et al., 2009), anti-TbBILBO1 antibody (1: 400 dilution) (Bonhivers et al., 2008), anti-TbGRASP polyclonal antibody (1:400 dilution) (He et al., 2004), anti-TbSAS-6 antibody (1:1,000 dilution) (Hu et al., 2015a), FITC-conjugated anti-HA monoclonal antibody (1:400 dilution, Sigma-Aldrich), and anti-Protein A polyclonal antibody (1:400 dilution, Sigma-Aldrich). Cells were washed with PBS and then incubated at room temperature for 1 h with the following secondary antibodies: FITC-conjugated anti-mouse IgG, Cy3-conjugated anti-rabbit IgG, Cy3-conjugated anti-mouse IgG, and FITC-conjugated anti-rabbit IgG, all of which were purchased from Sigma-Aldrich. Cells were washed with PBS for three times, mounted with DAPI-containing VectaShield mounting medium (Vector Labs), and imaged with the Olympus IX71 fluorescence microscope. Images were acquired and processed using the Slidebook software.

### Scanning electron microscopy

Scanning electron microscopy was performed as described previously (Souza Onofre et al., 2023). Cells were fixed with 2.5% (v/v) glutaraldehyde, and fixed cells were harvested by centrifugation at 750 ×*g* for 10 min, washed with PBS for three times, and then settled onto glass coverslips. Subsequently, cells on the coverslip were dehydrated in alcohol solutions (30%, 50%, 70%, 90%, and 100%) each for 5 min, and dried by critical point drying. Finally, cell samples were coated with a 5-nm metal film (Pt:Pd 80:20, Ted Pella Inc.) using a sputter-coater (Cressington Sputter Coater 208 HR, Ted Pella Inc.), and cells were imaged using Nova NanoSEM 230 (FEI) with the accelerating high voltage set at 8 kV and the scanning work distance set at 5 mm.

### Data analysis and statistical analysis

ImageJ (National Institutes of Health, Bethesda, MD; http://imagej.nih.gov/ij/) was used to measure the FAZ length, the flagellum length, the centrin arm length, the inter-centrin arm distance, the inter-FPC distance, and the inter-basal body distance, and to measure the fluorescence intensity of TbGRASP. Data were exported to Microsoft *Excel* or the GraphPad Prism9 for analysis. Statistical analysis was conducted using the two-tailed student’s t-test or one-way ANOVA (analysis of variance). Error bars represent standard deviation (SD) from the mean of three independent experiments.

## Acknowledgements

We thank Dr. Cynthia Y. He of National University of Singapore for providing anti-CC2D, anti-TbCentrin4, and anti-TbGRASP antibodies, Dr. Brooke Morriswood of University of Wurzburg for providing anti-TbMORN1 antibody, and Dr. Derrick Robinson of University of Bordeaux for providing anti-TbBILBO1 antibody. We are also grateful to Dr. Jianhua Gu of Houston Methodist Research Institute for assistance with scanning electron microscopy and to Dr. Li Li of the Clinical and Translational Service Center at University of Texas Health Science Center at Houston for assistance with mass spectrometry. This work was supported by the NIH R01 grants AI118736 and AI101437 to Z. L and in part by the Clinical and Translational Proteomics Service Center at the University of Texas Health Science Center at Houston. The content is solely the responsibility of the authors and does not necessarily represent the official views of the National Institutes of Health.

## Data availability

All data are contained within the manuscript.

## Author contributions

**Qing Zhou:** Conceptualization, Methodology, Visualization, Investigation, Formal analysis, Writing – Reviewing and Editing. **Huiqing Hu**: Methodology, Visualization, Investigation, Writing – Reviewing and Editing. **Ziyin Li**: Conceptualization, Supervision, Project administration, Funding acquisition, Writing – Original Draft, Writing – Reviewing and Editing.

---

**Supplemental Table S1. List of proximal proteins of KIN-G and WDR2 identified by BioID.**

**Supplemental Table S2. Mass spectrometry analysis of the phosphorylated KIN-G band pulled down by wild-type WDR2.**

**Supplemental Table S3. Mass spectrometry analysis of the non-phosphorylated KIN-G band pulled down by the WDR2-ΔMD mutant.**

**Supplemental Figure S1. Mass spectrometry identification of native KIN-G protein co-immunoprecipitated by 3HA-tagged wild-type WDR2 and the WDR2-ΔMD mutant.**

