## Supplemental Figure S1 for "WDR2 regulates the orphan kinesin KIN-G to promote hook complex and Golgi biogenesis in *Trypanosoma brucei*"

### Peptides of phosphorylated KIN-G pulled down by WT WDR2

MAKPLSGKAAPKNISVFLRVRPPVPRELKGG**T**FNNLVCDPSDPQRVTITRGGSARKGTKSFLFNRVFDPEC**T**QQTIYNEVARGAVDAAFDGQHGVLFVYG
QTGSGKTFTISNNDPEKPGVLQQSLRDIWDRFQADTEYDYSCTVSYVQLYNEMLTDLLDPQGGRVRIQLGPEGRGDVVLVTEASGASIERKVE**S**YEDCLK
YFYEGMDRKEMTSTKMNNTSSRSHTVFNFNLTRSAKVKTVDLSSAKANNEPVIALQGRLVVCDLAGSERASRTNAEGKTLDEATHINGSLLVLGKVVAAL
**T**ESGSQHAPFRESKLTRILQYSLLGNGNTSIVVNCSPCDDSTEETLGAIMFGQRAIQIKQDAKRHEILDYKALYYQLLADLDSKNDR**T**LETALSEERTAY
EDRIRVLEERIKIL**T**SENDMLRRESSQLGGTGPVSGTSTASGAAAAAVAMGGDDANDWRSLTMKMRRAIEKLDADLKRTDKERVELAQFLALEKNKVNVL
AQKLRAESLKHIMENKELTQRV**T**ELSIDNAKLKGTDYISFQPSAACEDALPLSLDSPRRGTPSSGLSQSINVGDAYLQEQLDKANRQLRVLNEERVELIV
YQMMASKAIRLLHAEK**T**SLANHLEKLKA

Phosphosites are underlined and bold, and are listed below:
Thr-32; Thr-72; Ser-194; Thr-301; Thr-388; Thr-415; Thr-523; Thr-617

### Peptides of non-phosphorylated KIN-G pulled down by WDR2-ΔMD

MAKPLSGKAAPKNISVFLRVRPPVPRELKGGTFNNLVCDPSDPQRVTITRGGSARKGTKSFLFNRVFDPECTQQTIYNEVARGAVDAAFDGQHGVLFVYG
QTGSGKTFTISNNDPEKPGVLQQSLRDIWDRFQADTEYDYSCTVSYVQLYNEMLTDLLDPQGGRVRIQLGPEGRGDVVLVTEASGASIERKVESYEDCLK
YFYEGMDRKEMTSTKMNNTSSRSHTVFNFNLTRSAKVKTVDLSSAKANNEPVIALQGRLVVCDLAGSERASRTNAEGKTLDEATHINGSLLVLGKVVAAL
TESGSQHAPFRESKLTRILQYSLLGNGNTSIVVNCSPCDDSTEETLGAIMFGQRAIQIKQDAKRHEILDYKALYYQLLADLDSKNDRTLETALSEERTAY
EDRIRVLEERIKILTSENDMLRRESSQLGGTGPVSGTSTASGAAAAAVAMGGDDANDWRSLTMKMRRAIEKLDADLKRTDKERVELAQFLALEKNKVNVL
AQKLRAESLKHIMENKELTQRVTELSIDNAKLKGTDYISFQPSAACEDALPLSLDSPRRGTPSSGLSQSINVGDAYLQEQLDKANRQLRVLNEERVELIV
YQMMASKAIRLLHAEKTSLANHLEKLKA
